# A conditional null allele of Dync1h1 enables targeted analyses of dynein roles in neuronal length sensing and neurological disorders

**DOI:** 10.1101/2022.02.20.481176

**Authors:** Agostina Di Pizio, Letizia Marvaldi, Marie-Christine Birling, Nataliya Okladnikov, Luc Dupuis, Mike Fainzilber, Ida Rishal

## Abstract

Size homeostasis is one of the most fundamental aspects of biology and it is particularly important for large cells as neurons. We have previously proposed a motor-dependent length-sensing and growth-regulating mechanism wherein a partial reduction in the levels of microtubule motor proteins should lead to accelerated neuronal growth. This prediction was originally validated in sensory neurons heterozygous for the Loa point mutation in dynein heavy chain 1 (*Dync1h1^Loa^*). Here we describe a new mouse model with a conditional allele allowing deletion of exons 24-25 in *Dync1h1*. Homozygous Islet1-Cre deletion of *Dync1h1* is embryonic lethal, but heterozygous animals (*Isl1-Dync1h1*^+/−^) survive to adulthood with approximately 50% dynein expression in targeted cell types. *Isl1-Dync1h1*^+/−^ adult sensory neurons reveal an accelerated growth phenotype, similar to that previously reported in *Dync1h1^Loa^* neurons. Moreover, *Isl1-Dync1h1*^+/−^ mice show mild impairments in gait, proprioception and tactile sensation; and slightly impaired recovery from peripheral nerve injury. Thus, conditional deletion of *Dync1h1* exons 24-25 enables targeted studies of the role of dynein in neuronal growth and neurological disorders.

## 1. Introduction

Neurons must extend their axons over long distances during development to reach their targets and establish functional circuits, and these extended neuronal arbors constitute a vulnerability prone to neurological diseases (Albus et al., 2013; Sleigh et al., 2019; Terenzio et al., 2017). A number of studies have described neurological disorders resulting from aberrations in signalling pathways implicated in neuronal size control (Nikolaeva et al., 2017; Sundberg and Sahin, 2020) or in intracellular transport (Sleigh et al., 2019). The dynein complex links intracellular transport with size control in neurons (Rishal and Fainzilber, 2019; Rishal et al., 2012), and indeed a number of studies have reported that dynein impairment underlies neurodegenerative diseases (Eschbach and Dupuis, 2011; Terenzio et al., 2018a). For example, a mouse point mutation in Dynein heavy chain 1 (*Dync1h1*), termed *Dync1h1^Loa^*, displays locomotor and motor system deficits arising from both motor and sensory neuron degeneration (Hafezparast et al., 2003; Schiavo et al., 2013).

We previously demonstrated that sensory neurons from *Dync1h1^Loa^* mice are characterized by increased axon length, in accordance with predictions from a model of a length-sensing mechanism based on motor-dependent bidirectional transport and the localization and local translation of key mRNAs in axons (Doron-Mandel et al., 2021; Perry et al., 2016; Rishal et al., 2012). However, interpretation of those results was complicated by the fact that the *Loa* mutation is a single nucleotide change rather than a clear loss of function allele. *Dync1h1^Loa^* mice and other described mutants are invariably non-viable as homozygotes, and the adult heterozygous animals have mild to severe neurodegeneration phenotypes (Terenzio et al., 2018a), raising the possibility that specific aspects of the phenotype might be due to effects of the point mutation rather than reduction in dynein levels. An allele deletion model would be of great utility to resolve such issues.

Here we describe a new conditional deletion allele for *Dync1h1* in the mouse. Islet1-Cre induced homozygous deletion mice are not viable, but heterozygosity is well tolerated, and mutant sensory neurons show the expected decrease in dynein levels, and increased axonal growth. Moreover, Islet1-Cre *Dync1h1*^+/−^ adult animals present deficits in motor and proprioception coordination, and delays in recovery from peripheral nerve injury. Thus, this new conditional *Dync1h1* allele enables targeted confirmation of dynein roles in neuronal growth control, and will facilitate comprehensive studies of the physiological roles of dynein.

## 2. Results

### 2.1 Generation of a conditional deletion allele for *Dync1h1*

We established a floxed *Dync1h1* allele targeting exons 24 and 25 of the *Dync1h1* gene, using available ES cell clones (Skarnes et al., 2011). The floxed allele was generated by transfection of the parental ES line with an Flp recombinase plasmid, leading to removal of the LacZ-NeoR cassette, and subsequent efficient germline transmission of the *Dync1h1* floxed allele. After recombination, exons 24 and 25 are deleted leading to a frameshift mutation prior to the motor domain of DYNC1H1 protein. This is the same targeting strategy recently used by Baehr and collaborators for analyses of *Dync1h1* roles in the retina and photoreceptor systems (Dahl et al., 2021a; Dahl et al., 2021b).

*Dync1h1* conditional exons 24-25 mice were bred with Islet1 (Isl1) Cre mice. The *Isl1* promoter drives expression in motor and sensory neurons (Srinivas et al., 2001), but also in heart and limb progenitors (Yang et al., 2006). Homozygous *Isl1-Cre* deletions of *Dync1h1* were embryonic lethal, but heterozygous mice (henceforth termed *Isl1-Dync1h1*^+/−^) survived to adulthood and were used for further characterization.

### 2.2 Reduced *Dync1h1* expression in heterozygous *Isl1-Dync1h1* mice

We first examined dynein mRNA and protein levels in dorsal root ganglion (DRG) neurons cultured from *Isl1-Dync1h1*^+/−^ mice. qPCR and Western blot analyses revealed a significant reduction of both *Dync1h1* mRNA **(Fig. 1A)** and protein **(Fig 1B)** in neurons cultured for 24 hr *in vitro*. This was further confirmed by immunostaining **(Figs 1C, D, and S1A, B)**. Proximity ligation assay (PLA) was then used to detect spatial coincidence of DYNC1H1 with Importin β1, a known component of dynein complexes in axoplasm and cytoplasm (Hanz et al., 2003; Perry et al., 2012). A clear decrease of axonal PLA signals for DYNC1H1/Importin β1 in heterozygous neurons confirms that reduced dynein expression is reflected in the prevalence of functional dynein complexes in the mutant **(Figs 1E, F, and S1C, D)**.

**Figure 1.**
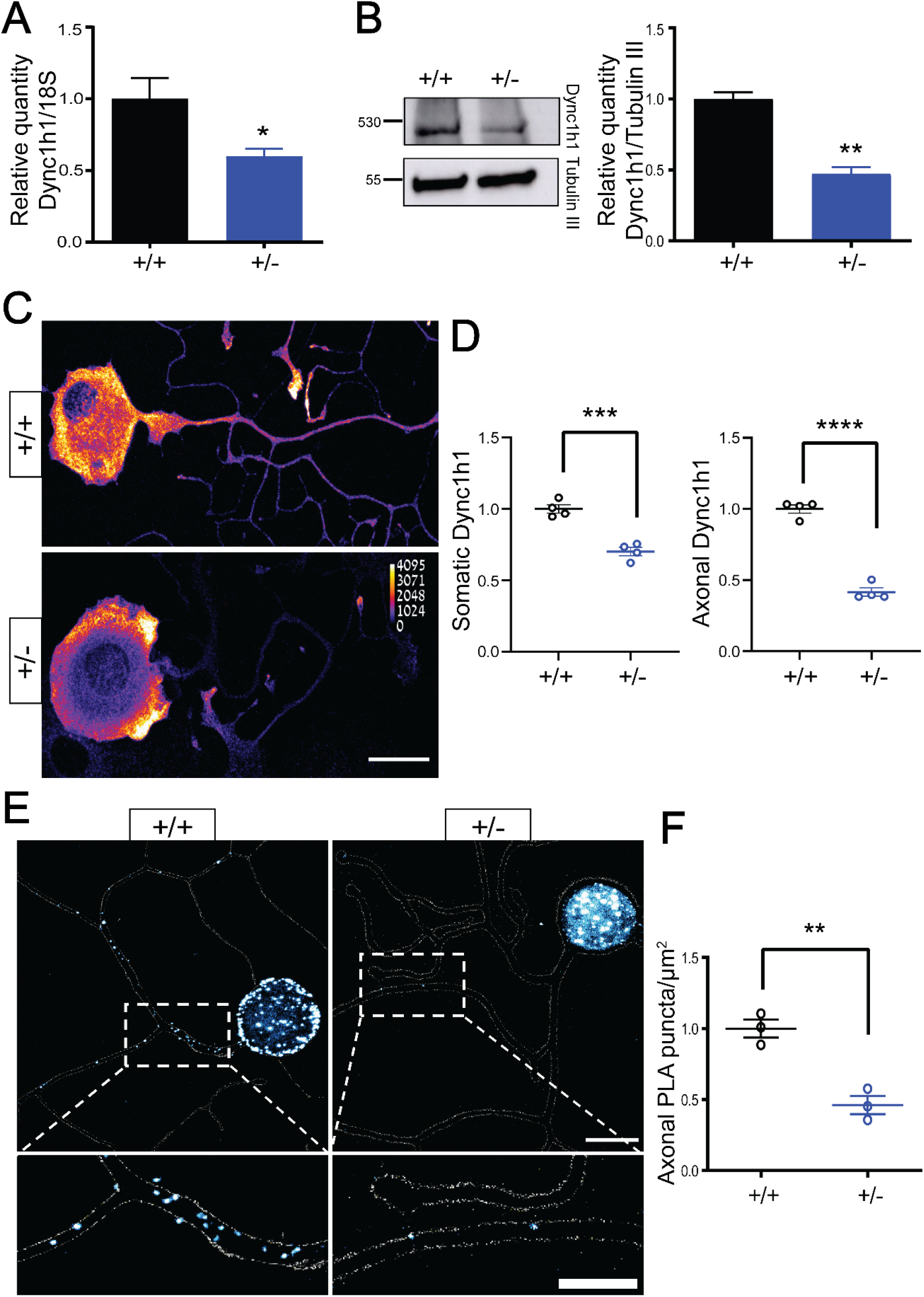
*Dync1h1* mRNA and protein levels are reduced in *Isl1-Dync1h1+/-* mice. **A,** qPCR quantification of Dync1h1 transcript using the ΔΔCt method on wild-type and heterozygous neurons 24 hr in culture, Mean ± SEM, N=4 biological repeats, * p < 0.05, unpaired t-test. **B,** Western blot analysis of lysates from wild-type and heterozygous 24 hr neuron cultures, Tubulin III served as a loading control, Mean ± SEM, N=3 biological repeats, ** p < 0.01, unpaired t-test. **C,** Representative images of immunostaining for Dync1h1 on wild-type and heterozygous neurons 24 hr in culture, scale bar 20 μm. **D,** Quantification of axonal and somatic immunoreactivity shows significant reduction of DYNH1C1 in mutant mice, Mean ± SEM, N=4 biological repeats, ***, **** represent p< 0.001, 0.0001, respectively, unpaired t-test. **E,** Proximity ligation assay (PLA) for DYNC1H1 and Importin β1 in wild-type and heterozygous neurons fixed after 24 hr in culture, scale bar 20 μm. Lower panels show magnification of the areas indicated by dashed borders, scale bar 10 μm. **F,** Quantification of the PLA shows reduction of axonal signal in *Dync1h1* mutant neurons, Mean ± SEM, N=3 biological repeats, ** p < 0.01, unpaired t-test.

### 2.3 Neuronal growth phenotype in heterozygous *Isl1-Dync1h1* cKO neurons

*Isl1-Dync1h1*^+/−^ mice presented abnormal hind limb posture when suspended by the tail **(Fig. S2)**, similar to the phenotype previously reported for whole body mutant *Dync1h1^Loa^* animals (Hafezparast et al., 2003), and indicative of motor and/or proprioceptive phenotypes (Chen et al., 2007). In previous work we had demonstrated enhanced neuronal growth of *Dync1h1^Loa^* sensory neurons, due to perturbation of a postulated dynein-dependent length-sensing mechanism (Rishal et al., 2012). We tested this prediction by cross-breeding *Isl1-Dync1h1*^+/−^ mice with Thy1-YFP mice (Feng et al., 2000) to obtain animals that express YFP primarily in large proprioceptive sensory neurons. Time-lapse imaging of axon outgrowth from YFP expressing neurons in culture reveal significantly accelerated growth of *Isl1-Dync1h1*^+/−^ genotype neurons, both at low culture densities that provide superior conditions for enhanced growth **(Fig. 2)**, and at higher densities comparable to the conditions used in previous studies on *Dync1h1^Loa^* mice **(Fig. S3)**.

**Figure 2.**
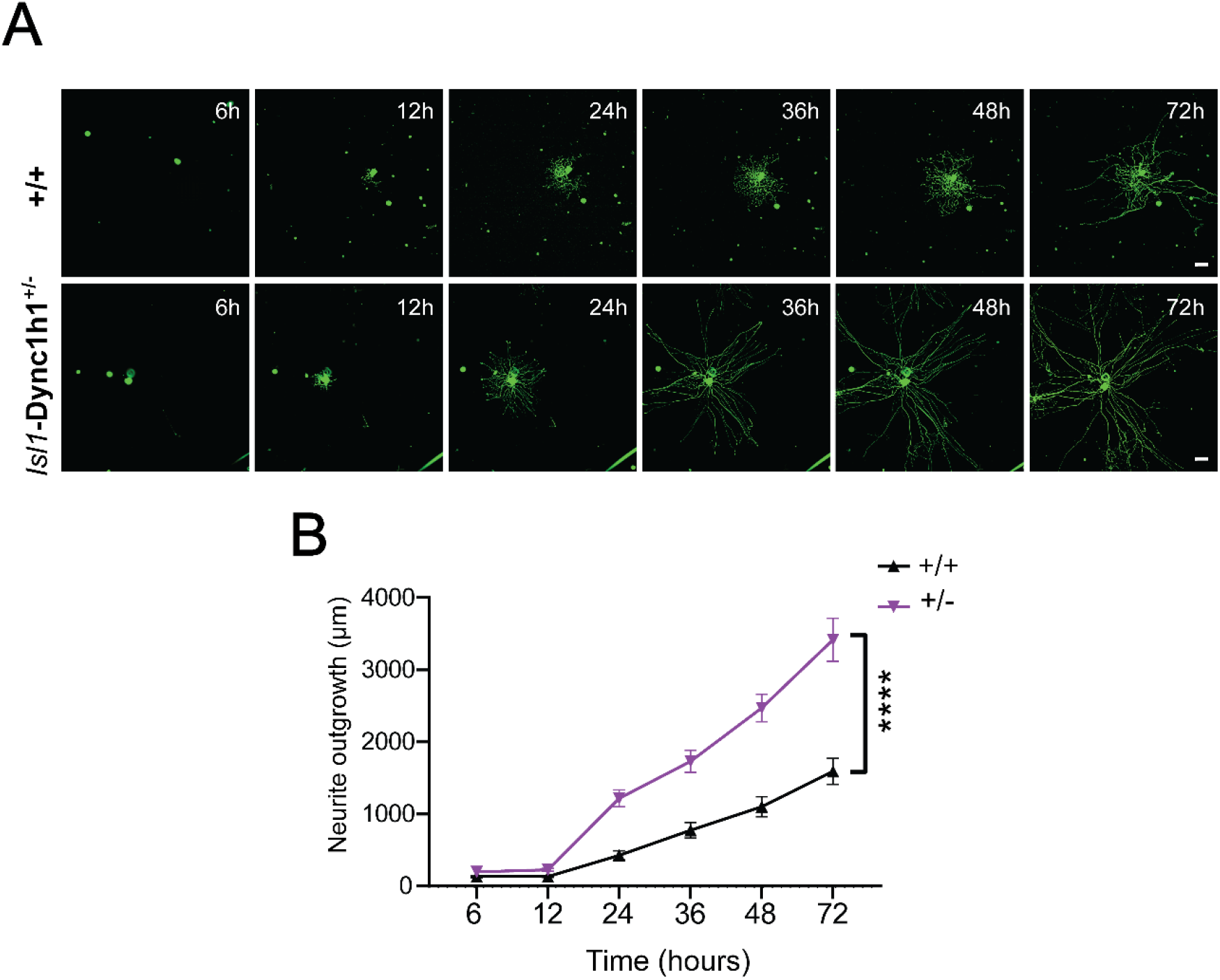
Time-lapse imaging reveals accelerated growth of *Isl1-Dync1h1+/-* sensory neurons. DRG neurons were plated at a density of approximately 25 YFP-expressing cells per well in 96-well plates and outgrowth was monitored by live cell imaging over 72 hr. **A,** Representative images of wild type and *Isl1-Dync1h1+/-* neurons at the indicated time points in culture. **B,** Quantification of neurite outgrowth, including only actively growing neurons in the analysis. The experiment was repeated four times. Mean ± SEM (n>315 growing neurons from four biological repeats, **** p<0.0001, two way ANOVA). Scale bar 100 µm.

### 2.4 *Isl1-Dync1h1*^+/−^ mice reveal deficits in motor coordination, gait and injury responses

*Dync1h1^Loa^* mice and other dynein heavy chain heterozygous point mutant animals have clear behavioral deficits, including abnormal hind limb positioning and decreased grip strength (Terenzio et al., 2018a). We evaluated locomotion deficits, impaired balance or muscular weakness in *Isl1-Dync1h1*^+/−^ animals, using catwalk and rotarod tests (Dumont, 2011). *Isl1-Dync1h1*^+/−^ mice revealed mild but significant impairments in both assays **(Fig. 3A, B)**. Alterations in tactile sensitivity were also observed **(Fig. 3C)**, while in contrast there were no deficits in heat sensitivity **(Fig. 3D)**.

**Fig. 3.**
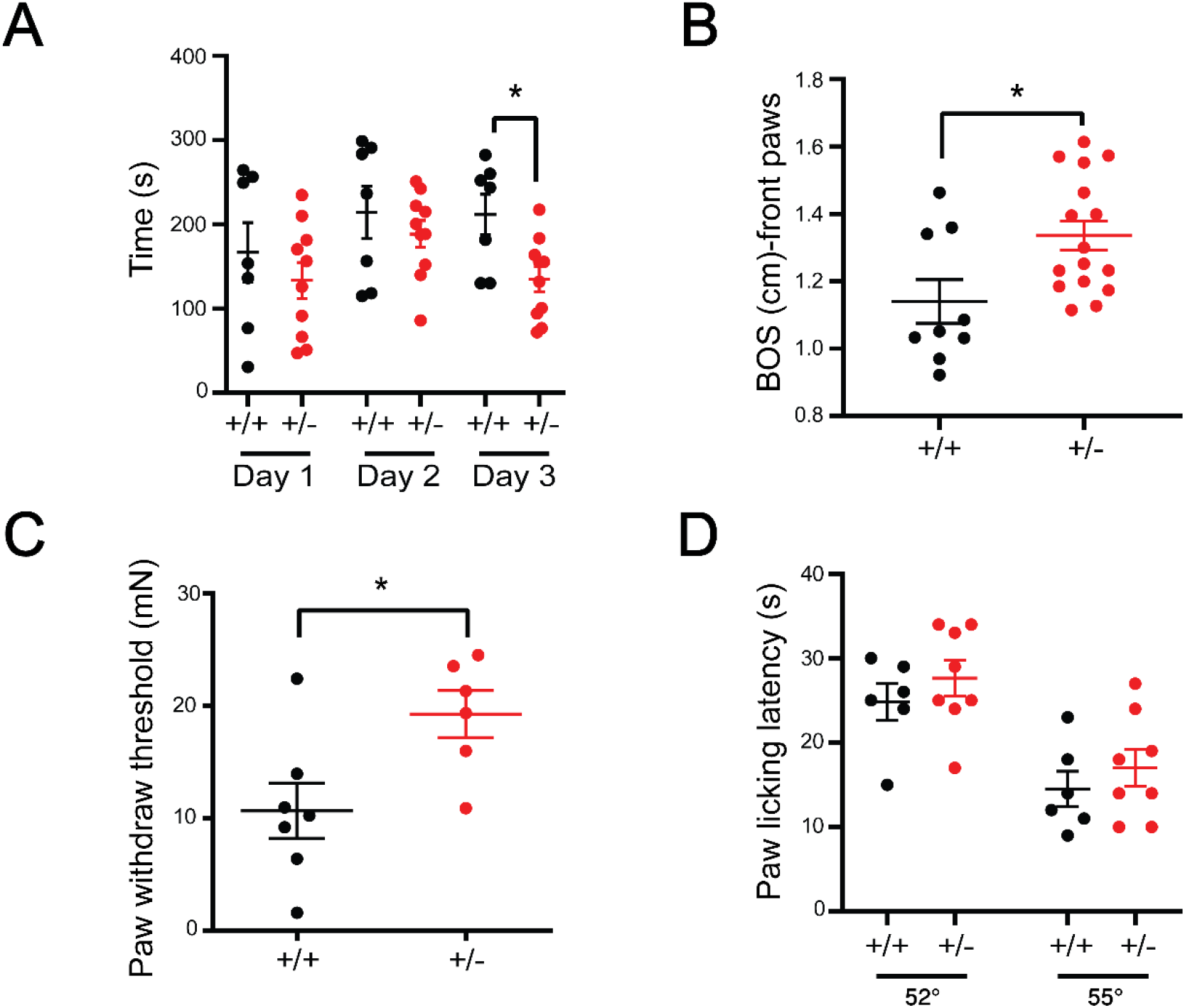
Motor coordination, gait and tactile sensitivity are impaired in heterozygous *Islet1*-Dync1h1 cKO mice, while no defects in thermal sensitivity were identified. **A,** Rotarod test. Mice underwent 3 days of training and were subjected to 4 trials each day. The rotarod was accelerated from 0 to 40 rpm in 4 min for the first two days and from 0 to 40 rpm in 2 min the third day. Latency to fall was registered and the average of trials of each day shows significantly reduced performance of heterozygous mice during the third day. Mean ± SEM (n>9, * p < 0.05, two way ANOVA). **B,** Walking gait was assessed by Catwalk analysis, revealing wider base of support in the mutant mice. Mean ± SEM (n>9, * p < 0.05, unpaired t-test). **C,** Von Frey tests show reduced mechanical sensitivity of *Isl1-Dync1h1*^+/−^ animals. Mean ± SEM (n>6, * p < 0.05, unpaired t-test). **D,** Hot plate tests at 52oC or 55oC show no differences in thermal sensitivity.

Previous studies have shown a critical role for dynein in retrograde signaling after peripheral nerve injury (Rishal and Fainzilber, 2014; Terenzio et al., 2017). We therefore examined injury responses of sensory neurons in *Isl1-Dync1h1*^+/−^ mice. Sciatic nerve conditioning lesion (Smith and Skene, 1997) enhances elongating growth of wild type sensory neurons cultured from L3/L5 DRG, but there was no such effect on *Isl1-Dync1h1*^+/−^ neurons **(Fig. 4)**. The dynein deficient neurons exhibit enhanced growth in culture under basal conditions **(Figs 2** and **S3)**, and this may mask any potential effect of the conditioning lesion. We then proceeded to examine functional recovery from sciatic nerve lesion in *Isl1-Dync1h1*^+/−^ mice *in vivo*. Mice gait parameters were evaluated on a catwalk apparatus before crush injury of the sciatic nerve, and over a time course up to 28 days post-lesion (**Fig. 5**). As expected, both wild type and *Isl1-Dync1h1*^+/−^ mice reduce usage of the injured limb immediately after injury, with gradual recovery of different gait parameters over time after lesion (**Fig. 5**). Differences between wild type and *Isl1-Dync1h1*^+/−^ mice were apparent through the recovery time course, with *Isl1-Dync1h1*^+/−^ mice showing somewhat reduced recovery of the injured limb at the time points assayed during the recovery phase (**Fig. 5B-D**).

**Fig. 4.**
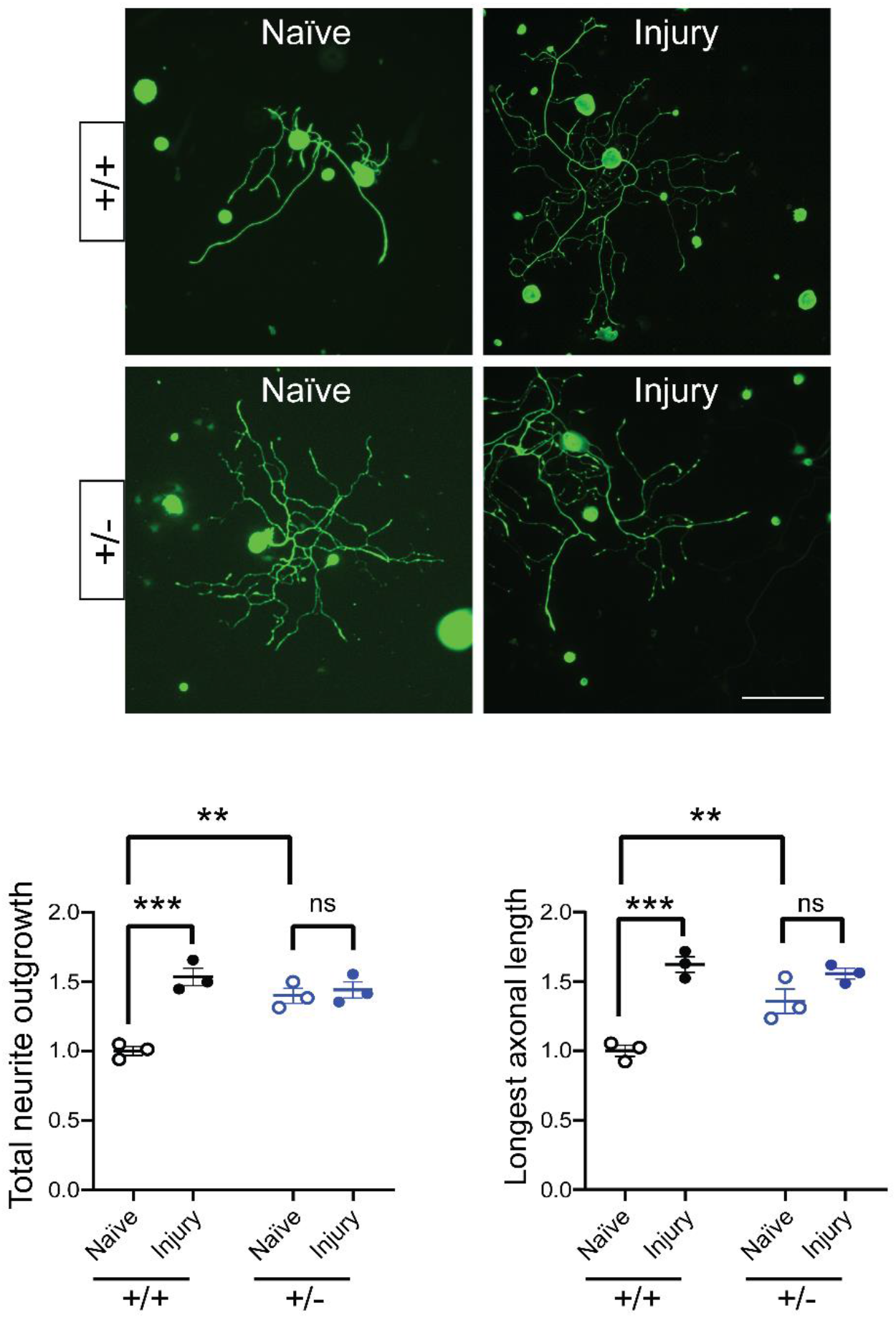
Conditioning lesion enhances axonal outgrowth in wild-type neurons but not in *Dync1h1* mutant neurons. Wild-type and heterozygous mice were subjected to unilateral sciatic nerve crush. 3 days after injury, L3, L4 and L5 DRGs were dissociated and cultured for 24 hr. Quantification of total neurite outgrowth and longest axonal length show that *Dync1h1* mutant neurons grow longer than the wild-type in naive condition, but don’t respond to the conditioning lesion. Mean ± SEM (N=3 biological repeats, **, *** p < 0.01, 0.001, one way ANOVA). Scale bar 200 µm.

**Fig. 5.**
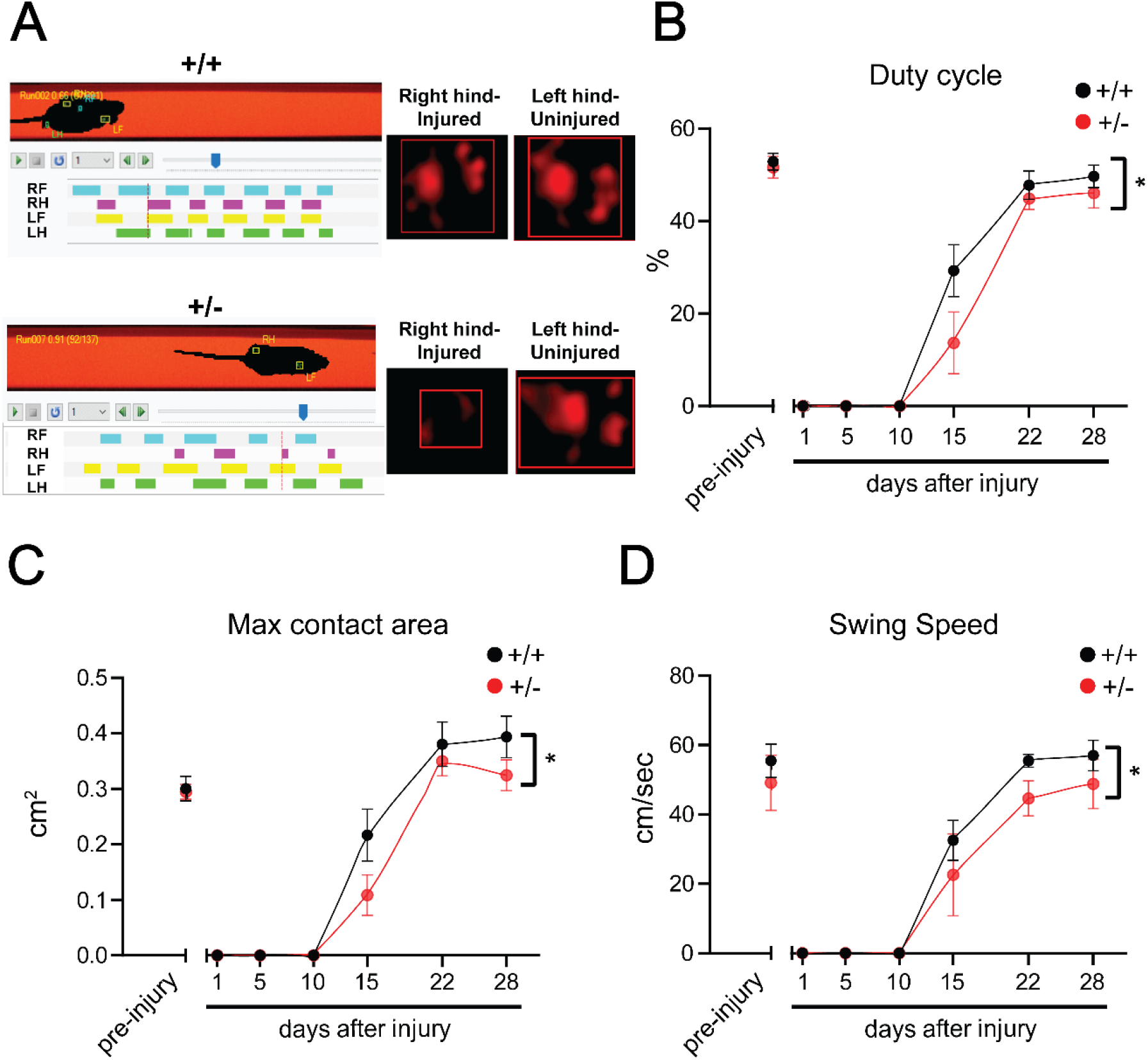
Gait analysis reveals somewhat slower regeneration of *Isl1-Dync1h1+/-* mice. **A,** Representative images of catwalk gait analysis at 15 days after sciatic nerve crush at the right hind (RH) leg. *Isl1-Dync1h1+/-* mice have slower recovery of the injured leg (B-D). **B,** Duty cycle: duration of the contact of the paw with the glass plate, expressed as percentage of a step cycle. **C,** Max contact area: surface area of the print at the point of maximum contact. **D,** Swing speed: speed (cm/sec) of the paw during swing (phase of no contact with the glass plate). Mean ± SEM (n>7, * p < 0.05).

## 3 Discussion

The dynein heavy chain *Dync1h1* is an essential gene in multicellular organisms, and accordingly all constitutive *Dync1h1* mouse mutants described to date are embryonic lethal when homozygous (Terenzio et al., 2018a). Heterozygous mutants are mostly viable, albeit typically with progressively worsening neurodegeneration phenotypes, and since the mutation is organism-wide the origin of a specific deficit may be unclear. For example, a number of studies in both *Dync1h1^Loa^* and *Dync1h1^Cra^* mice suggest that their originally described motor phenotypes may be a secondary consequence of earlier effects in sensory neurons (Chen et al., 2007; Dupuis et al., 2009; Ilieva et al., 2008). Alleles allowing cell-type and/or temporal specificity in targeting dynein will be of value in resolving such issues. Moreover, potential confounding or masking effects are less likely in a deletion allele as compared to an expressed mutant with altered amino acid sequence. Indeed, a recent study reported photoreceptor maintenance phenotypes upon conditional deletion of *Dync1h1* exons 24 and 25 in retinal cells (Dahl et al., 2021a), that were not reported previously in *Dync1h1* point mutant mice.

The *Dync1h1* allele described herein enables Cre-driven deletion of dynein with the expected effects on dynein expression and function in targeted cell types and tissues. Our previous work had postulated roles for dynein in retrograde injury signalling (Rishal and Fainzilber, 2014) and in intrinsic length-sensing and regulation of axonal growth rates (Rishal et al., 2012). The length-sensing model predicts axon length increase upon partial reduction of dynein levels (Rishal and Fainzilber, 2019; Rishal et al., 2012), and initial experiments in sensory neurons of *Dync1h1^Loa^* mice supported this prediction (Rishal et al., 2012). The results reported above in *Isl1-Dync1h1*^+/−^ sensory neurons further support the model, with clearly enhanced outgrowth of heterozygous dynein knockout neurons. Thus, the *Isl1-Dync1h1*^+/−^ model phenocopies *Dync1h1^Loa^* effects on sensory neuron growth *in vitro*, and on limb proprioception *in vivo*. Taken together, these findings suggest that the sensory and motor phenotypes observed in the *Dync1h1^Loa^* mutant are caused by loss of dynein function, rather than a dominant-negative effect of mutant *Loa* DYNC1H1 (Garrett et al., 2014; Ori-McKenney et al., 2010).

*In* vivo characterization of Isl1-Dync1h1^+/−^ mice reveal further parallels with Dync1h1^*Loa*^ in limb coordination (Hafezparast et al., 2003), and altered proprioception and gait (Ilieva et al., 2008). The Isl1-Dync1h1^+/−^ mice also show changed tactile, but not thermal, sensitivity, indicating physiological impacts of dynein reduction in different sensory neuron subtypes. As regards injury response, Isl1-Dync1h1^+/−^ neurons do not exhibit enhanced growth after conditioning lesion, but this might be due to masking of the injury response due to their basal accelerated growth phenotype. However, Isl1-Dync1h1^+/−^ mice show somewhat impaired recovery after nerve injury *in vivo*, indicating that the enhanced basal neuronal growth of dynein heterozygote neurons is not sufficient to overcome other deficiencies caused by the deletion.

Taken together, our results on heterozygous *Dync1h1* deletion in sensory neurons confirm model predictions and previous findings from *Dync1h1^Loa^* mice on a dynein-dependent mechanism that regulates neuronal growth. Importantly, a number of rare neurological diseases have been genetically associated with mutations in the *DYNC1H1* gene. These diseases include motor neuron diseases such as spinal muscular atrophy with lower extremity predominance (SMA-LED) or Charcot Marie Tooth diseases (Becker et al., 2020; Harms et al., 2012; Scoto et al., 2015; Weedon et al., 2011), but also neurodevelopmental defects (Poirier et al., 2013) and/or epilepsy (Lin et al., 2017; Matsumoto et al., 2021). The availability of the dynein deletion allele model will be helpful to establish cell specific consequences of dynein loss in the context of these diseases with widely divergent clinical phenotypes.

## 4 Materials and methods

### Animals

All animal experiments were approved by the IACUC (Animal Care and Use Committee) at the Weizmann Institute of Science. Mice were bred and maintained at the Veterinary Resources Department in a room with 12 hr light-dark cycle. Experiments were carried out on animals between 2-6 months old.

### Mutant mouse line establishment

The *Dync1h1* conditional mutant mouse line was established at the Institut Clinique de la Souris - PHENOMIN (http://www.phenomin.fr). Three clones were ordered from the Mutant Mouse Resource Research centers (https://www.mmrrc.org/), and validated by Southern blot with an internal Neo probe (Codner et al., 2021). The presence of the 3’ LoxP site was confirmed by PCR. Clones were karyotyped by chromosome spreading and one selected clone was electroporated with a plasmid expressing the Flp recombinase to remove the flipped LacZ-NeoR cassette and obtain the conditional knock-out allele (tm1c). PCR confirmed the excision of the flipped cassette. One positive clone was microinjected in blastocysts, male chimeras were obtained and germ line transmission was achieved (tm1c allele; conditional knock-out).

### Cultures and Antibodies

DRG culture preparations were performed as previously described (Terenzio et al., 2018b). Primary antibodies used in this study are the following: chicken anti-NFH (Abcam, ab72996, 1:2000), rabbit anti-dync1h1 (Proteintech, 12345-1-AP, 1:500), mouse anti-Importin β1 (monoclonal, generated in-house), rabbit anti-mTOR (Cell Signaling, #2972, 1:1000), rabbit anti-Nucleolin (Abcam, ab50279, 1:500) and rabbit anti β-III tubulin (abcam, ab18207, 1:2000). Secondary antibodies used for immunostaining are anti-chicken/mouse/rabbit/ Alexa Fluor 488/594/647 (Jackson Immunoresearch, 1:1000). Secondaries for immunoblots were HRP-conjugated rabbit antibodies (Bio-Rad Laboratories, 1:10000).

### Dync1h1 gene expression analysis by qPCR

The mRNA level of Dync1h1 was quantified by q-PCR analysis. RNA was extracted from DRG neurons grown 24h in culture using the Oligotex mRNA Mini kit (Quiagen). Superscript III (Invitrogen) was used to synthesize cDNA while qPCR reactions were prepared in a total volume of 20ul using PerfeCTa SYBR Green (Quanta Biosciences, Gaithersburg, USA) and performed on a ViiA7 System (Applied Biosystem). 18S was used as housekeeping gene for normalization and data were analysed using the comparative ∆∆Ct method (Livak and Schmittgen, 2001). The sequences of the primers are the following:

18S forward: AAACGGCTACCACATCCAAG
18S reverse CCTCCAATGGATCCTCGTTA
Dync1h1 forward: CCAACAGCTTGGCGTTCAT
Dync1h1 reverse: GGGACGACACTGGCTTGTCT

### Dync1h1 protein expression analysis by Western blot

Wester blot analysis was conducted on DRG neurons cultured for 24h. Cells were harvested and lysed in RIPA buffer. Proteins were then boiled in 5x Laemmli sample buffer, fractionated by SDS-PAGE, and transferred to 4-15% gradient gels using a Bio-Rad transfer apparatus according to the manufacturer’s protocol. Membranes were incubated for 1 hr at room temperature in a blocking solution of 5% milk in TBST (10 mM Tris, pH 8.0, 150 mM NaCl, 0.5% Tween 20), washed and incubated with the rabbit anti-dync1h1 antibody and rabbit anti-β-III tubulin antibody for overnight at 4°C. The following day, they were washed and incubated with HRP-conjugated anti-rabbit antibodies for 1 hr. Images were captured with the ECL system (Amersham Biosciences) and the signal quantified using *Fiji* software.

### PLA labelling

Proximity ligation assay (PLA) (tom Dieck et al., 2015) was used to detect spatial coincidence of Dync1h1 and Importin β1 proteins. PLA was performed according to manufacturer’s instructions using Duolink (Sigma: PLA probe anti-mouse minus DUO92004, anti-rabbit plus DUO92002, and detection kit red DUO92008). After the PLA protocol, cells were immunostained with chicken anti-NFH antibody. The PLA signal was quantified with the analysed particle function of Fiji software, using a mask based on NFH staining and dividing the number of PLA positive puncta by the area.

### Imaging

Images of neuronal for outgrowth were taken at 10x magnification on an ImageXpress Micro Confocal (Molecular Devices). Imaging for immunostaining was performed at 60x on Olympus confocal microscopes, either a FV100 UPLSAPO-NA 1.35 (oil-immersion objective) or a Fluoview FV10i (water-immersion objective).

### Behavioral profiling

All assays were performed under dim illumination (~10 lx) during the “dark” active phase of the diurnal cycle. Mice were tested with the ROTOR-ROD, Catwalk, von Frey and Hot plate tests as follows.

#### ROTOR-ROD

mice underwent 3 days of training on the rotarod, accelerating from 0 to 40 rpm in 4 min (inclination of 10 rpm/min) during the day 1 and 2. On the third day, the rotarod was accelerated from 0 to 40 in 2 min (20rpm/min). Each day, mice were subjected to 4 trials with a 2 min break in between as shown previously (Ezra-Nevo et al., 2018). Latency to fall (s) was recorded and the average of the 4 trials per day was considered.

#### Catwalk

the test was carried on as previously described (Terenzio et al., 2020). Motivation was achieved by placing the home cage at the runway end and each mouse was tested 3-5 times. The collected data were analyzed using the Catwalk Ethovision XT10.6 software (Noldus Information Technology, The Netherlands). Mice showed differences in the base of support parameter (BOS), defined as the average width (cm) between the paws during each step cycle.

#### Von Frey

in this test, mechanical sensitivity was tested by pressing filaments of different diameters on the plantar surface of the mouse’s paw. Before starting the test, mice were habituated in chambers suspended above the test apparatus wire mesh grid for an hour. Once the mice were calm, the test with the Von Frey filaments started and the response was considered positive if the paw was sharply withdrawn upon filament application, starting with 13.7 millinewton filaments and then progressing in an up-down method, as previously described (Marvaldi et al., 2020).

#### Hot plate

Mice were tested for heat sensitivity at temperatures of 52° and 55°. Each mouse was placed in a 20 cm high Plexiglas box on the heated metal surface and the latency to initiate a nociceptive response (licking, paws shaking, jumping) was recorded.

### Conditioning lesion

For conditioning lesion experiments, mice were subjected to unilateral sciatic nerve injury and, after three days, DRGs L3, L4, L5 were dissociated and cultured for 24h. The plating was done on a DMEM/F12 medium supplemented with N1 and 10% fetal bovine serum, as was done previously (Hanz et al., 2003). Neurons were then fixed in 4% paraformaldehyde, stained with NFH, and subjected to morphological analysis. Images were analyzed using the Metamorph software (Molecular Devices).

### In vivo regeneration

To test regeneration in vivo, we made use of the Catwalk test described above. Before sciatic nerve injury, mice were trained on the Catwalk apparatus and a baseline of both genotypes was performed one day before the crush. The locomotion recovery of the mice was observed during the day 1, 5, 10, 15, 22 and 28 post-injury showing reduced recovery of the Isl1-Dync1h1^+/−^ mice.

## 5. Acknowledgements

We thank Marco Terenzio, Dalia Gordon, Vladimir Kiss and Michael Tsoory for excellent professional expertise and helpful advice, and Jérôme Sinniger, Nir Sharabi and Hagit Dafni for technical support. This work was funded by Agence Nationale de la Recherche, France (ANR-10-JCJC-1101 to LD), the Motor Neuron Disease Association (Dupuis/Apr16/852-791 to LD), and the Israel Science Foundation (ISF 1337/18 to MF and IR). MF is the incumbent of the Chaya Professorial Chair in Molecular Neuroscience at the Weizmann Institute of Science.

---

**Figure S1.**
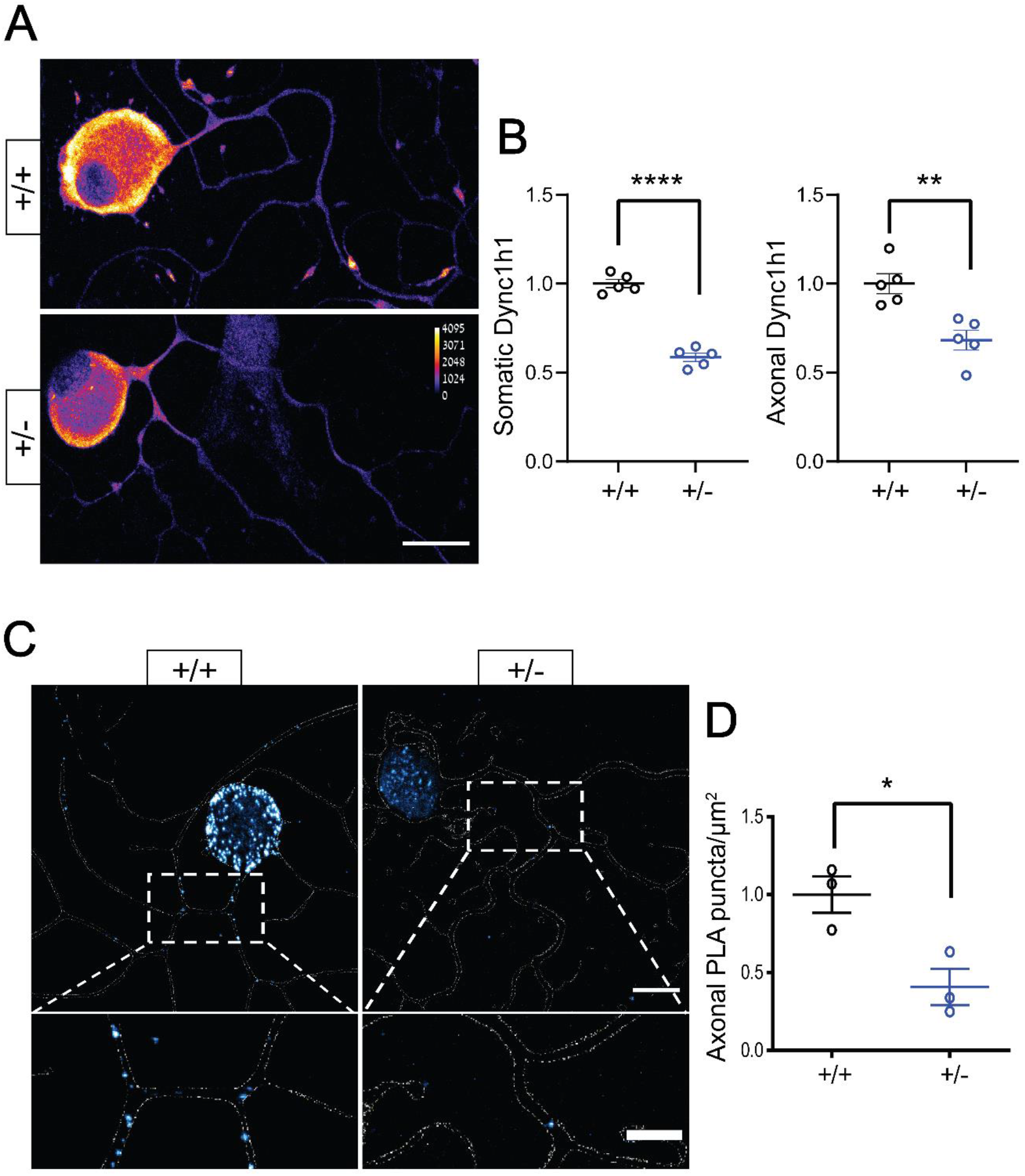
Dync1h1 protein levels are reduced in *Isl1-Dync1h1*^+/−^ mice 48 hr in culture. **A,** DYNC1H1 immunostaining on wild-type and heterozygous neurons 48 hr in culture, scale bar 20 μm. **B,** Quantification of (A), Mean ± SEM, N=5 biological repeats, **, **** p< 0.01, 0.0001, respectively, unpaired t-test. **C,** PLA for DYNC1H1 and Importin β1, Scale bars 20 μm upper panels, 10 μm lower magnified views. **D,** PLA quantification, Mean ± SEM, N=3 biological repeats, * p< 0.05, unpaired t-test.

**Figure S2.**
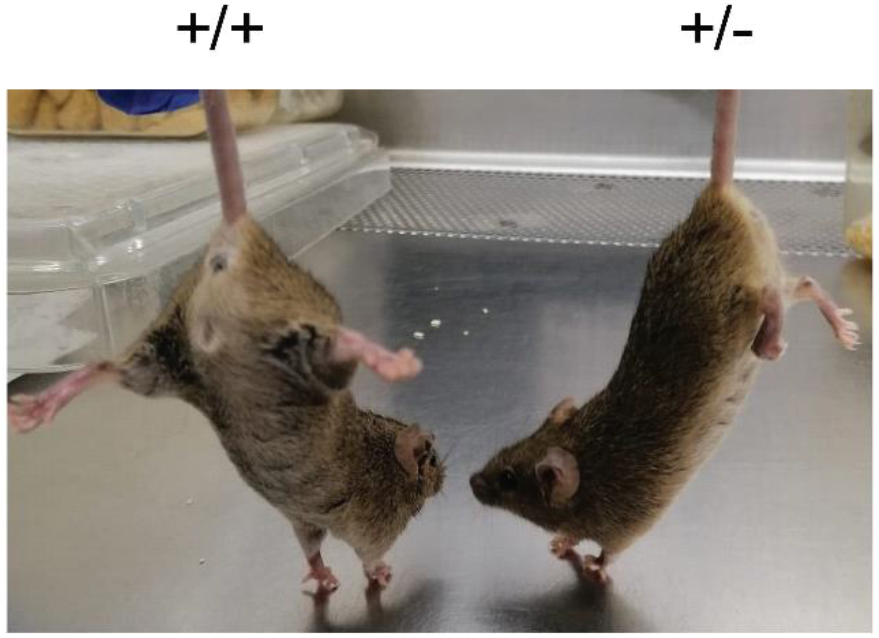
*Isl1-Dync1h1*^+/−^ mice present an abnormal clenched hind limb posture when suspended by the tail, resembling the characteristic “legs at odd angles” phenotype of *Dync1h1^Loa^* mice.

**Figure S3.**
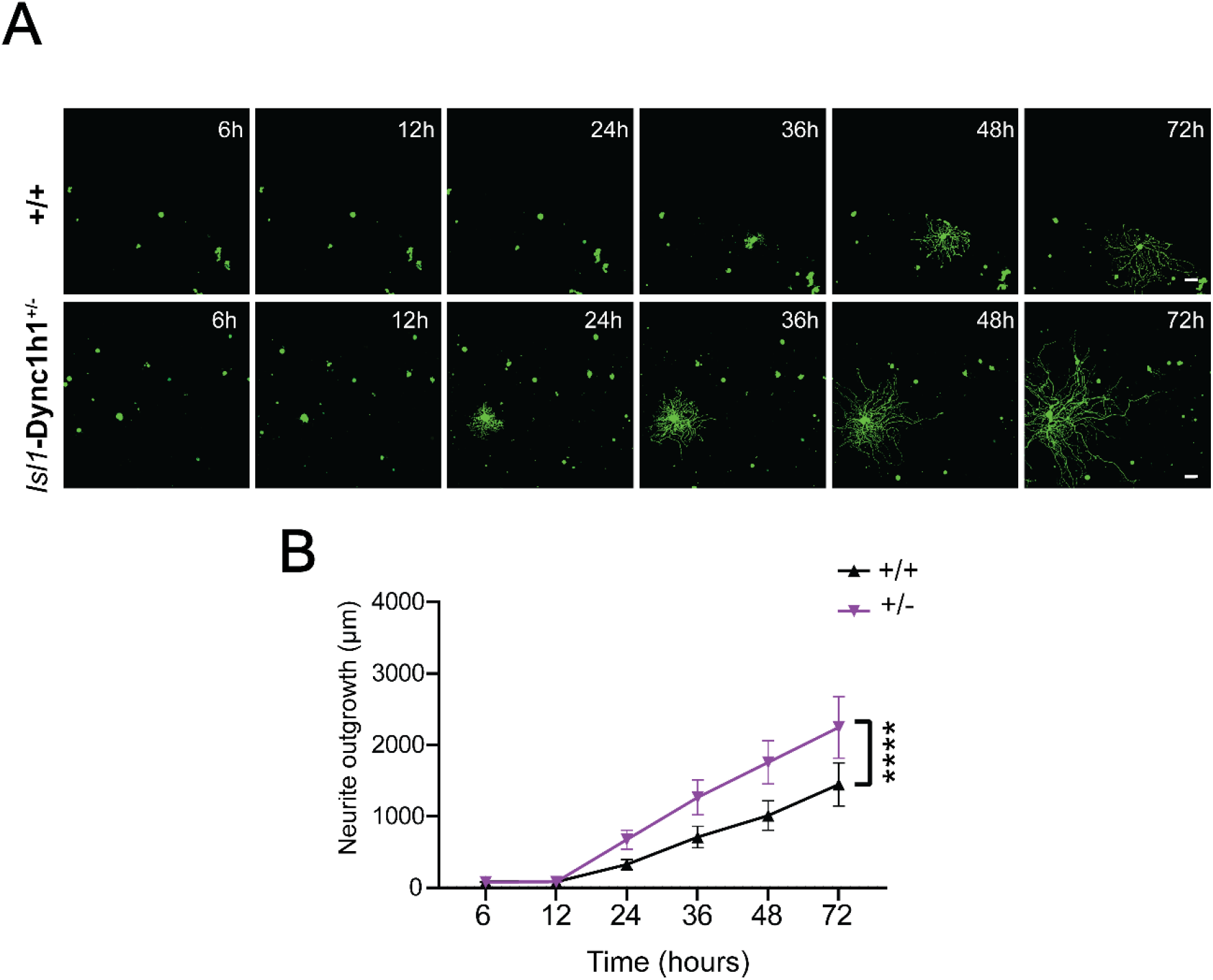
Time-lapse imaging reveals accelerated growth of *Isl1-Dync1h1^+/−^* sensory neurons also when plated at higher density. Experimental conditions as in Figure 2, except for plating density which was 45 YFP-expressing neurons per well in 96-well format. **A,** Representative images of wild type and *Isl1-Dync1h1^+/−^* neurons at the indicated time points in culture. **B,** Quantification of neurite outgrowth, including only actively growing neurons in the analysis. Mean ± SEM (n> 128 growing neurons, **** p<0.0001, two way ANOVA). Scale bars 100 µm.

## Notes

### Competing Interest Statement

The authors have declared no competing interest.

